# Heritage of ancient cultures supports conservation: a continent-wide perspective from the Eurasian steppes

**DOI:** 10.1101/2022.09.29.510067

**Authors:** Balázs Deák, Ádám Bede, Zoltán Rádai, Iwona Dembicz, Iva Apostolova, Péter Batáry, Róbert Gallé, Csaba Albert Tóth, József Dózsai, Ivan I. Moysiyenko, Barbara Sudnik-Wójcikowska, Georgi Nekhrizov, Fedor N. Lisetskii, Zhanna A. Buryak, Szabolcs Kis, Sándor Borza, Laura Godó, Tatyana M. Bragina, Ilya Smelansky, Ábel Molnár, Miklós Bán, Ferenc Báthori, Zoltán Árgay, János Dani, Orsolya Valkó

## Abstract

Civilisations including ancient ones, have shaped the global ecosystems in many ways through a co-evolution of landscapes and humans. However, the cultural legacies of ancient and lost civilisations are seldom considered in conservation. Here using a continental-scale dataset containing over 1,000 data records on the localities, land cover, protection status and cultural values related to ancient steppic burial mounds (so-called ‘kurgans’), we evaluated how these iconic and widespread landmarks can contribute to grassland conservation in the Eurasian steppes, which is one of the most endangered biomes on Earth. By using Bayesian logistic generalized regressions and proportional odds logistic regressions, we aimed to reveal the potential of mounds in preserving grasslands considering landscapes with different levels of land use transformation. We also compared the conservation potential of mounds situated inside and outside protected areas and assessed whether the presence of cultural, historical or spiritual values support the maintenance of grasslands on them. We revealed that kurgans have enormous importance in preserving grasslands in transformed landscapes outside protected areas, where they can act as habitat islands, and provide an additional pillar for conservation by contributing to habitat conservation and improvement of habitat connectivity. We found that besides their steep slopes hindering ploughing, the existence of cultural, historical or religious values could almost double the chance for grassland occurrence on kurgans due to the related extensive land use and the respect of local communities. As the estimated number of steppic mounds is about 600,000 and also similar historical features exist in all continents, our results can be upscaled to a global level. Our results also suggest that an integrative socio-ecological approach in conservation might support the positive synergistic effects of conservational, landscape and cultural values.

## Introduction

Grasslands covering 26% of the Earth’s terrestrial areas, have a crucial role in agricultural production and provision of essential ecosystem services (Dixon et al., 2014; Terreret al., 2021). The vast Eurasian steppe represents the largest proportion of global temperate grasslands (>10 million km^2^) (Kirschner et al., 2020; Wesche et al., 2016). Steppe grasslands formed under continental climate and dry habitat conditions harbour high biodiversity and an enormous number of endemic plant and animal species (Chytry et al., 2022; Kirschner et al., 2020). The extended grasslands of the Eurasian steppe belt provided proper habitats for large herds of migrating herbivores in prehistoric times and have later been used as pastures for livestock, determining the everyday life and lifestyle of local human populations. Pastoral communities and their grazing herds became an integral part of the open landscapes for millennia and formed the habitat structure and species composition of vast areas (Ventresca Miller et al, 2020).

Civilisations including ancient ones, have shaped the global ecosystems in many ways through a co-evolution of landscapes and humans. However, the cultural legacies of ancient and lost civilisations are seldom considered in conservation. During the Late Copper and Early Bronze Age (3100–2500 BC), the extensive grasslands of the Pontic-Caspian steppes witnessed the rise of ancient nomadic herders, the Yamnaya culture, which profoundly shaped the history of Europe (Wilkin et al., 2021). Due to many technological advances, such as the domestication of the horse, the combination of horse traction and bulk wagon transport and the diet change towards meat and dairy consumption, this prehistoric pastoralist community was able to expand its home range by thousands of kilometres along the Eurasian steppes, which provided suitable environmental conditions for its nomadic herder lifestyle (Anthony & Brown, 2011; Wilkin et al., 2021). As shown by ancient DNA evidence, the Yamnaya expansion extended over a 6,000 km wide area reaching Central Europe in the West and the Altai Mountains in the East (Allentoft et al., 2015; Haak et al., 2015). By fundamentally transforming the European genetic landscape and introducing the basics of the currently spoken Indo-European tongues, the Yamnaya expansion even at a distance of 5,000 years still has a considerable effect on today’s Eurasian societies (Haak et al., 2015). More importantly, the Yamnaya Culture also had a lasting effect on the landscape as its burial mounds still are the most widespread man-made prehistoric structures in the Eurasian steppe landscape (Deák et al., 2016).

Ancient burial mounds (also named kurgans) typical to the Eurasian steppes are distributed from Hungary to Mongolia. Although their original number was presumably higher by an order of magnitude (millions of kurgans were destroyed during the past centuries due to agricultural intensification and infrastructural development), there are still about 400,000– 600,000 existing kurgans in the steppes (Deák et al., 2016). An overwhelming proportion of kurgans were built by the Yamnayas, but both preceding and subsequent steppe cultures also built kurgans during the Iron Age and in the Migration Period (Gimbutas, 2000). Originally kurgans had religious function as they served as burial sites and sacred places, which were visible from large distances in the vast open steppes. Most kurgans were created by piling soil on top of a pit grave that was sunk under the soil surface^13^. Their diameter ranges from a few meters up to 100 meters, and they are usually 0.5–15 meters high. Besides steppe kurgans, there are ten thousands of burial mounds with similar appearance and cultural functions both in Europe (e.g. Czech Republic Hejcman et al., 2013; Germany: Dreibrodt et al., 2009; England: Andrews & Fernandez-Jalvo, 2012; Denmark: Andersen, 2012) and in North America (USA: Steponaitis, 1986).

Recent ecological studies revealed that millennia-old kurgans often hold dry grasslands and can act as habitat islands in transformed landscapes providing refuge for grassland species and maintaining high biodiversity (Apostolovaetal., 2022; Deák et al., 2020; Dembicz et al., 2020; Sudnik-Wójcikowska et al., 2011). All studies agree that the potential of kurgans for preserving grasslands is connected with their steep slopes, which impede ploughing (Deák et al., 2020; Dembicz et al., 2020). High environmental heterogeneity caused by the steep slopes of kurgans facing different cardinal directions also supports the co-existence of various microhabitats from mild to harsh even within a small area (Deák et al., 2021a; Lisetskii et al., 2016). By harboring grassland specialist species with different habitat requirements, kurgans are often characterized by a higher plant diversity compared to plain grassland areas of the same size (Deák et al., 2021a). Although it has not been directly studied yet, the cultural values of kurgans also seem to support the maintenance of grassland habitats.

The biodiversity potential of kurgans was demonstrated by regional surveys from Ukraine (Sudnik-Wójcikowska et al., 2011), Hungary (Deák et al., 2020) and Bulgaria (Apostolova et al., 2022)(Fig. 1). Despite the small overall area of the surveyed kurgans (UA: 106 kurgans, 100.7 ha; HU: 138 kurgans, 33.6 ha; BG: 111 kurgans, 32.8 ha), they hold an enormously high plant biodiversity (UA: 721; HU: 469; BG: 1,059 species) representing a considerable proportion of the three countries’ floras (UA: 14%; HU 17%; BG: 26%) with a high number of rare and protected species (UA: 71; HU: 73; BG: 45 species). Although studies on animal biodiversity preserved by the kurgans are scarce, data from Hungary suggest that kurgans are also important sites for animal conservation. Among red-listed species, 21 ant, 18 orthopteran, 76 true bug and 20 rove beetle species were recorded in 138 surveyed kurgans (Deák et al., 2020). Besides being biodiversity hotspots in agricultural landscapes, kurgans can also increase landscape-scale plant diversity in vast pristine steppes by harboring unique species confined exclusively to the special environmental conditions provided by the kurgans (Deák et al., 2017).

**Fig. 1.**
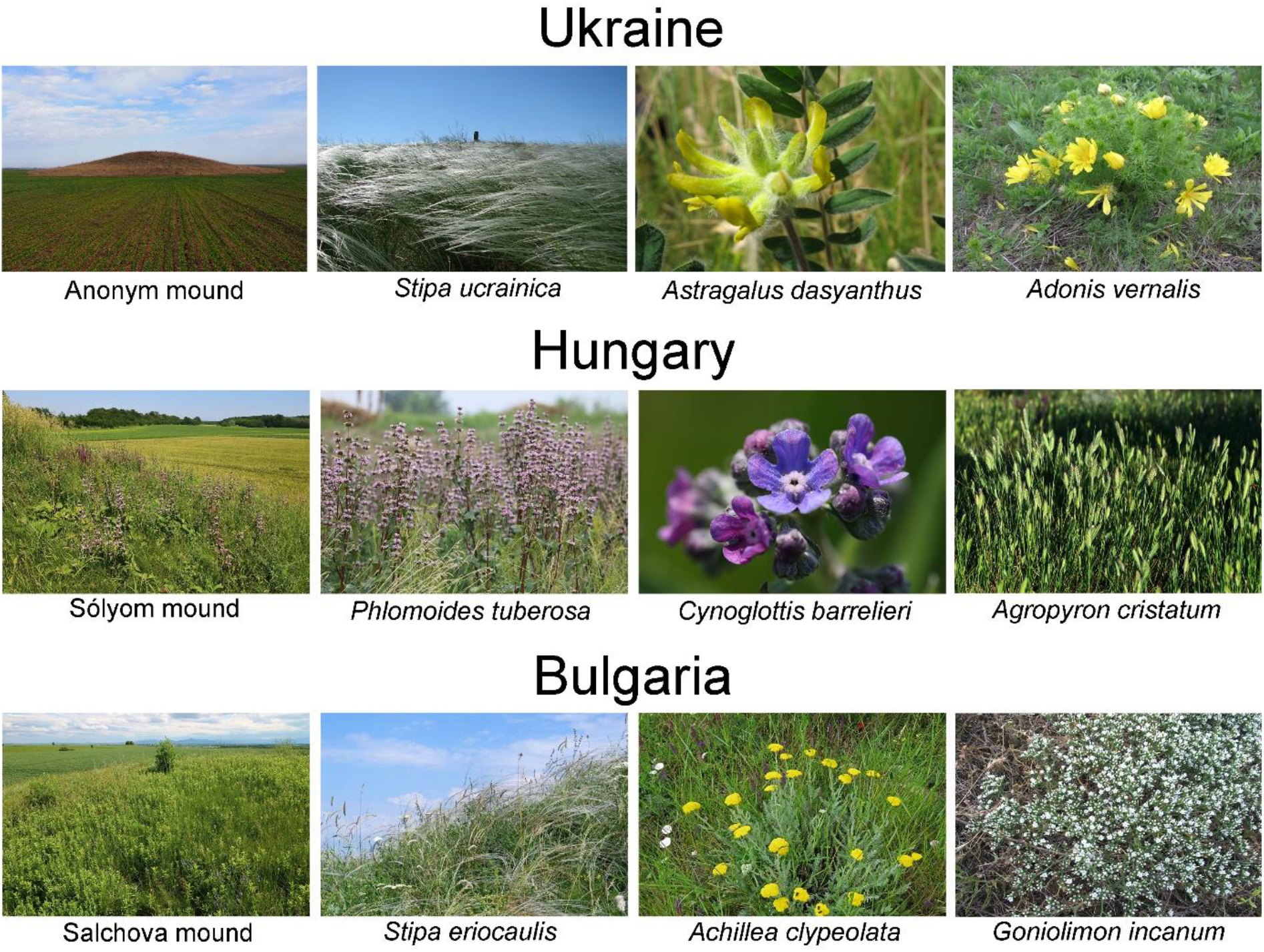
Examples of kurgans from Ukraine, Hungary and Bulgaria of kurgans preserving species-rich grassland vegetation and typical red-listed species confined to them. Photo credits: I. I. Moysiyenko (1–4); B. Deák (5–9), I. Apostolova (10–12).

Due to the globally increasing human pressure on grassland ecosystems, the steppes in which kurgans are embedded are exposed to serious threats. Both their area and conservation status have been declined for centuries in an alarming and constant rate (Dixon et al. 2014; Wesche et al., 2016). The protection of steppe ecosystems is especially challenging as overall coverage and mean individual size of protected areas (PAs) is the lowest in this biome (Kirschner et al., 2020; Wesche et al., 2016). Similar to other grassland ecosystems, the most serious threat to steppes is the habitat loss due to conversion into crop fields or forest plantations and the expansion of urban infrastructure (Biró et al., 2018; Kamp et al., 2016). The last remnants of large and near-natural steppes have been included into the network of PAs in the western part of the steppe biome during the past decades. However, in intensively used lowland landscapes, the coverage of the PAs is in general extremely low compared to mountainous regions because of the high population density and competing economic interests in available fertile land (Bhagwat & Rutte, 2006).

Under such settings in densely populated and intensively used agricultural landscapes, remaining small grassland fragments might be of outstanding importance, but their conservation poses great challenges due to their smallness and scattered distribution (Dudley et al., 2009, Maxwell et al., 2020). Still, grassland fragments situated outside PAs have a crucial role in the maintenance of biodiversity and grassland-related ecosystem services and also in providing functional spatial connections between meta-populations of grassland biota (Deák et al., 2020; Maxwell et al., 2020). A considerable proportion of the Eurasian kurgans hold such grassland fragments, but their conservation potential is insufficiently explored on a continental scale.

Despite the high cultural and conservation importance of kurgans, there is a huge lack of knowledge on their actual number, localities, conservation status and cultural values in most countries of the steppe biome. Databases containing information relevant for conservation management or monitoring purposes have been missing in most of the countries so far. We filled this knowledge gap by developing an open access, up-to-date geo-referenced kurgan inventory, which provides a comprehensive overview of the locations and characteristics of kurgans at continental level (Fig. 2). The authors of this paper founded and maintain the Eurasian Kurgan Database (EKDB; http://openbiomaps.org/projects/kurgan (Deák, 2019a; Deák et al., 2019b), which contains essential information on the conservation status of kurgans covering a wide geographic range including understudied regions (Deák et al., 2019b). In this paper we provide a continental-level overview of the role of kurgans in grassland conservation by using data assembled in the EKDB. We aimed to reveal whether kurgans situated outside PAs have conservation relevance by preserving grasslands. We also studied the correlation between cultural values and conservation potential using four types of protection (non-protected, area-based, cultural and combined protection; please see in Methods).

**Fig. 2.**
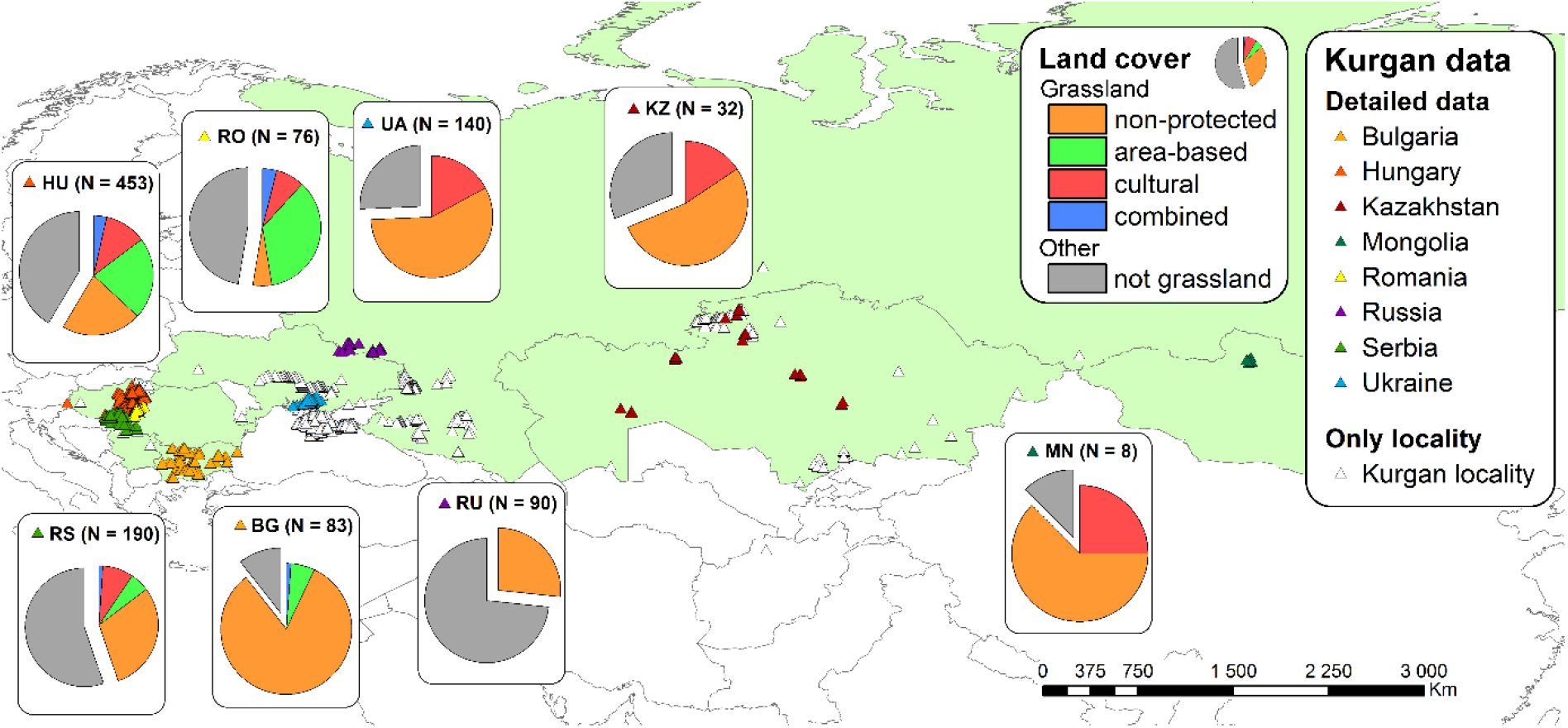
Map of the kurgans registered in the Eurasian Kurgan Database. The number of kurgans from which detailed data were available is indicated in parentheses after the abbreviation of the country name (n=1072). Pie charts represent the proportion of kurgans that does not hold grasslands and the share of the four protection types (non-protected, area-based, cultural and combined protection) within the kurgans covered by grasslands. Map lines delineate study areas and do not necessarily depict accepted national boundaries.

## Methods

### Kurgan data

For our study we used kurgan data that was collected and uploaded to the Eurasian Kurgan Database (EKDB; http://openbiomaps.org/projects/kurgan; Deák, 2019a) by the authors of this paper. The EKDB, which uses an open-source database framework (OpenBioMaps; Bán et al., 2022), is a public online database providing interfaces for uploading and accessing kurgan-related data for a wide spectrum of users. Since the EKDB was developed as a citizen science tool, the attributes in the data forms were selected considering that data providers are not exclusively biologists or geographers. Hence the necessary data can be reliably recognized or estimated without any previous professional training and can be recorded by using the GPS and the camera of a mobile phone. This allows to cover a large geographical region and to collect a high number of records. Besides the name and the geographical position of a kurgan, more detailed data on the physical attributes, the landscape context, the conservational status and the cultural, historical and religious values can be collected in the EKDB. The database also allows photos of the surveyed kurgans to be added. Currently the EKDB contains 3,808 records (detailed data on 1,072 kurgans and further 2,736 kurgan localities).

In the present study we evaluated only the records with detailed data (Deák et al., 2022); kurgan localities were only used for upscaling our results. The surveyed kurgans were situated in eight countries harboring zonal or extrazonal steppes or forest-steppes. Although we used the largest available dataset on kurgans, there are still many underrepresented regions such as Kazakhstan and Mongolia. Existing literature (Bourgeois & Gheyle, 2007; Deák et al., 2016) suggests the presence of a high number of kurgans in these countries, but they are still unexplored. Each record was validated using satellite images provided by Quantum GIS (QGIS Development Team) and the photos available in the EKDB.

In the study we focused on the presence of grassland on the kurgans, the amount of grassland in the surrounding landscape, and presence of kurgan-related cultural values (Table 1). We categorized the kurgans into four groups based on their protection status and presence/absence of cultural values: (i) ‘non-protected’: situated outside PAs, without cultural values; (ii) ‘area-based protection’: situated inside PAs, without cultural values; (iii) ‘cultural protection’: situated outside PAs, but holding cultural values; (iv) ‘combined protection’: situated inside PAs and holding cultural values.

**Table 1.**
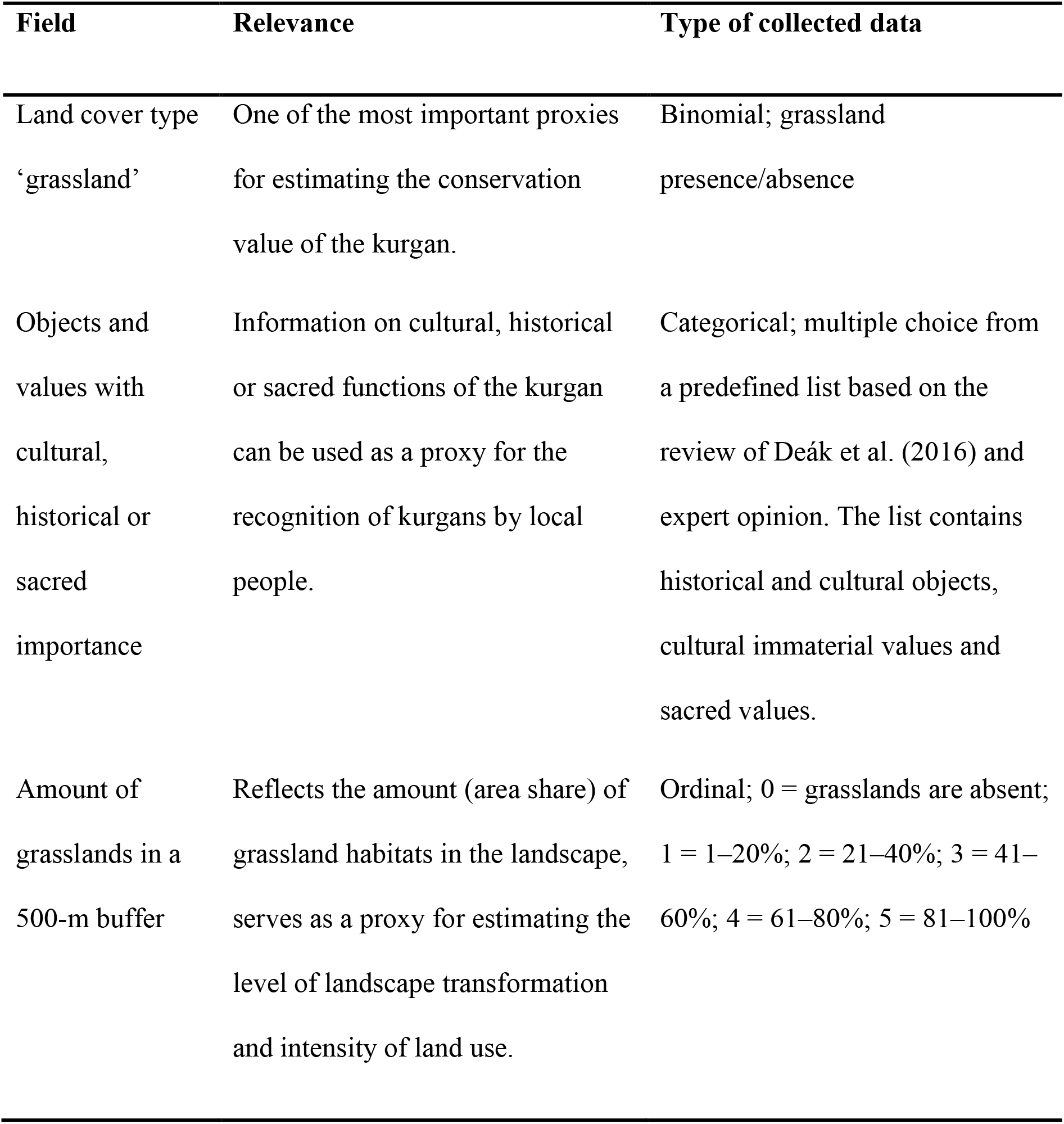
Kurgan attributes used for the analyses (source: Eurasian Kurgan Database).

| Field | Relevance | Type of collected data |
| --- | --- | --- |
| Land cover type<br>'grassland' | One of the most important proxies for estimating the conservation value of the kurgan. | Binomial; grassland presence/absence |
| Objects and values with cultural, historical or sacred importance | Information on cultural, historical or sacred functions of the kurgan can be used as a proxy for the recognition of kurgans by local people. | Categorical; multiple choice from a predefined list based on the review of Deák et al. (2016) and expert opinion. The list contains historical and cultural objects, cultural immaterial values and sacred values. |
| Amount of grasslands in a 500-m buffer | Reflects the amount (area share) of grassland habitats in the landscape, serves as a proxy for estimating the level of landscape transformation and intensity of land use. | Ordinal; 0 = grasslands are absent; 1 = 1–20%; 2 = 21–40%; 3 = 41–60%; 4 = 61–80%; 5 = 81–100% |

We used the thematic layers of the World Database on Protected Areas (WDPA, 2022) for identifying kurgans situated in protected areas. We considered all kinds of protected areas listed in the WDPA, such as areas under federal protection and Natura 2000 sites, except Special Areas of Conservation since these areas are primarily designated for bird conservation and do not necessarily focus on preserving natural or semi-natural habitats.

In Hungary kurgans have a special protection status, i.e. they are protected by the Nature Conservation Law (Act LIII. of 1996 on Nature Conservation; NCL). This protection is not area-based, but prohibits any activities that might threaten the landscape-related or conservation values of the kurgans. In addition, most of the kurgans are considered “protected landscape elements” in the Hungarian regulations concerning the single area payment scheme (SAPS) of the European Union’s Common Agricultural Policy (CAP). Farmers who apply for an area-based subsidy cannot cultivate the kurgans or take any actions (e.g. ploughing or afforestation) that would endanger their integrity. This particular type of protection was not considered in the analyses, i.e. Hungarian kurgans were, like those from other countries, assigned to the four protection categories listed above.

### Statistical analyses

All data handling and statistical analyses were carried out in R (R Core Team). For exploring the effect of kurgan protection category (non-protected, area-based protection, cultural protection, combined protection), grassland amount in the neighboring landscape and country (explanatory variables) on grassland presence on the kurgan (quasi binomial response variable), we used Bayesian logistic generalized regression models with the R-package “arm” (Gelman & Su; 2020). For logistic generalized regression models, we used Bayesian GLMs because they enable the estimation of reasonable standard errors (SE) even for low-variability or invariant group levels. We used the “quasi-binomial” model family in the logistic regression models to control for imbalanced data distribution caused by the differences in the amount of kurgan data from different countries. For exploring the differences in grassland amount in the surrounding landscape (ordinal response variable) among kurgan protection categories and countries, we used proportional odds logistic regression models with the R-package “MASS”. Given the low number of kurgan data reported from Kazakhstan (32) and Mongolia (8), we excluded these countries from the statistical analyses, but considered them in the Discussion. Due to the imbalanced data distribution across countries, we specified weights based on the proportion of observations originating from different countries in all models. Specifically, we used inverse probability weighting (w*i* = 1 – pC*i*, where w*i* is the weight for the *i*th observation and pC*i* is the proportion of observations originating from the country C to which the *i*th observation belongs) to control for disparate sample sizes from the different countries. To avoid multicollinearity (correlations between explanatory variables) and biases from imbalanced co-occurrences of levels of multiple categorical explanatory variables, we fitted separate models on each factor. Estimated marginal means and contrasts from ordinal models, mean probabilities and odds-ratios from logistic models were acquired using the “emmeans” R-package (Lenth, 2019). We adjusted p-values from the fitted models with Bonferroni’s method to decrease the probability of type I errors. In the Results we report estimated marginal means (EMMs) and contrasts for ordinal models and estimated mean probabilities and odds ratios for binomial models.

## Results

We found that 58% of the kurgans were covered by grasslands. Kurgans situated in Russia held grasslands with the lowest (27.0%), while Bulgarian (89.8%), and Ukrainian (74.5%) kurgans with the highest probability. In Hungary (59.3%), Romania (51.8%) and Serbia (44.9%) probability of grassland occurrence on kurgans was intermediate (Fig. 3.; Appendices S1-3). Landscape surrounding the kurgans was highly transformed in Russia and Ukraine. Grassland amount in the landscape was the highest in Romania and Bulgaria and was intermediate in Hungary and Serbia (Fig. 4.; Appendices S4-6).

**Fig. 3.**
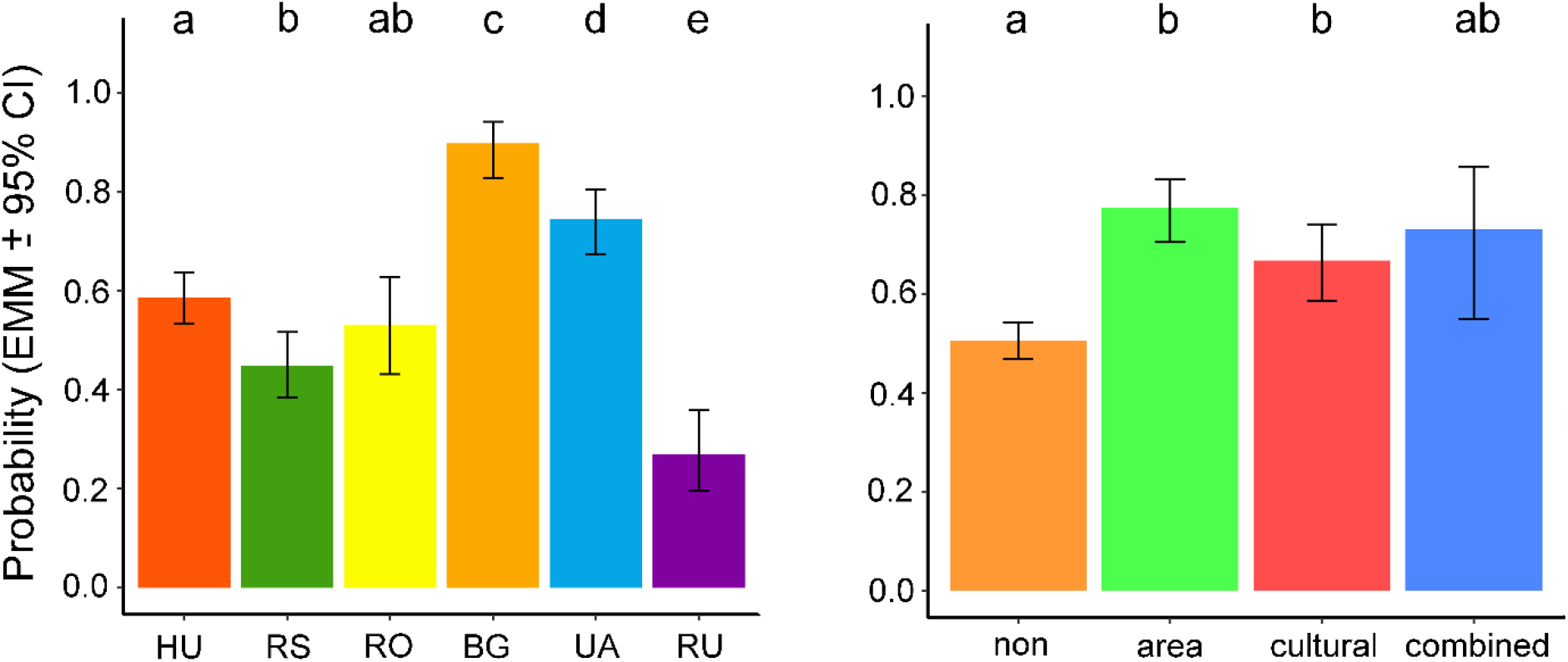
Estimated marginal mean (EMM) probability of grassland occurrence on kurgans situated in the six studied countries (on the left) and on kurgans with different protection types (on the right). Different letters denote significant differences between groups (contrasts are from Bayesian logistic generalized regression models; p≤0.05) (n=1032).

**Fig. 4.**
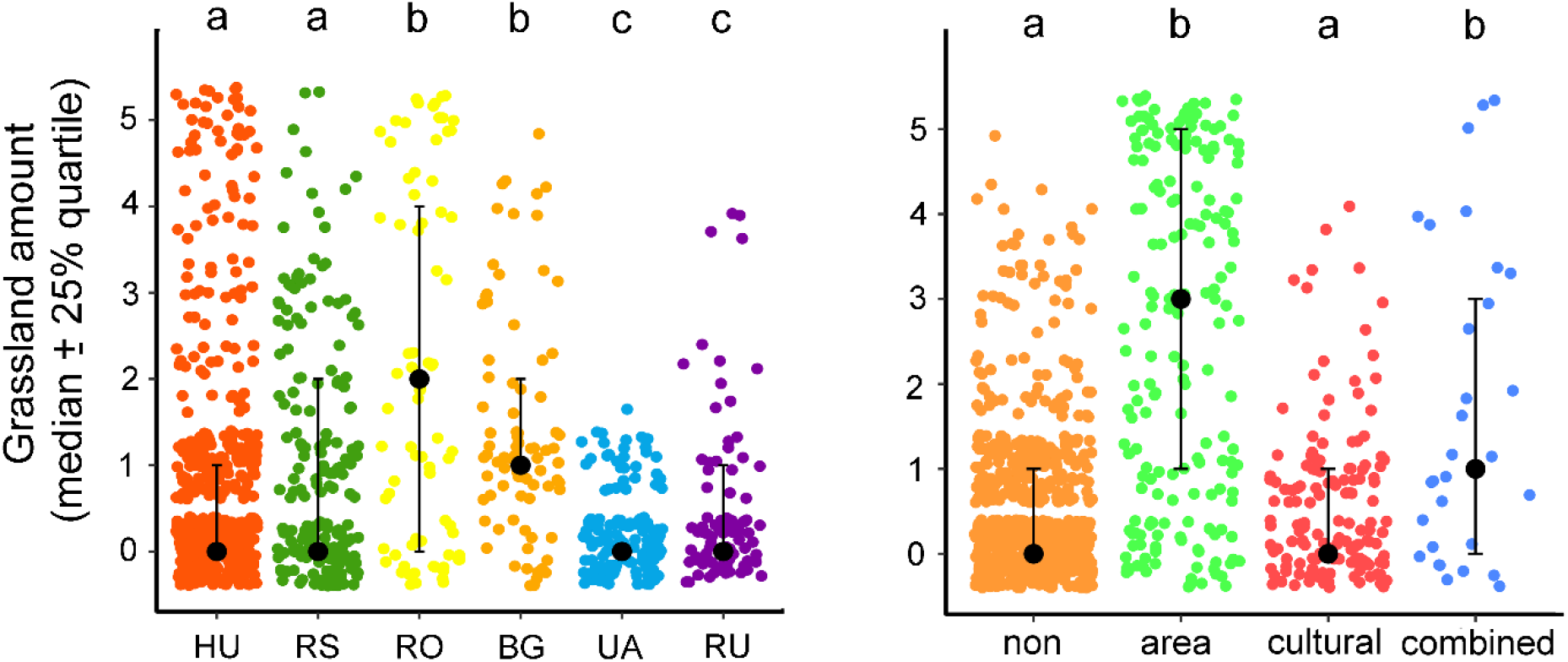
Grassland amount in the landscape (within 500 m buffer) around kurgans situated in the six studied countries (on the left), and around kurgans with different protection types (on the right). Grassland amount is represented on an ordinal scale: 0 = grasslands are absent; 1 = 1–20%; 2 = 21–40%; 3 = 41–60%; 4 = 61–80%; 5 = 81–100%. Different letters denote significant differences between groups (contrasts are from proportional odds logistic regression model; p≤0.05) (n=1032).

In landscapes where grasslands were absent, the probability of grassland presence on kurgans was the lowest (39%) (Fig. 5.; Appendices S7–9). In landscapes where the grassland amount in the neighboring landscape ranged between 1 and 40%, the probability of grassland presence on the kurgans was almost three times higher (odds ratio 0.31), and reached up to 71%. In landscapes with a grassland share of >40%, the probability for grassland presence on kurgans considerably increased and was the highest (>95%).

**Fig. 5.**
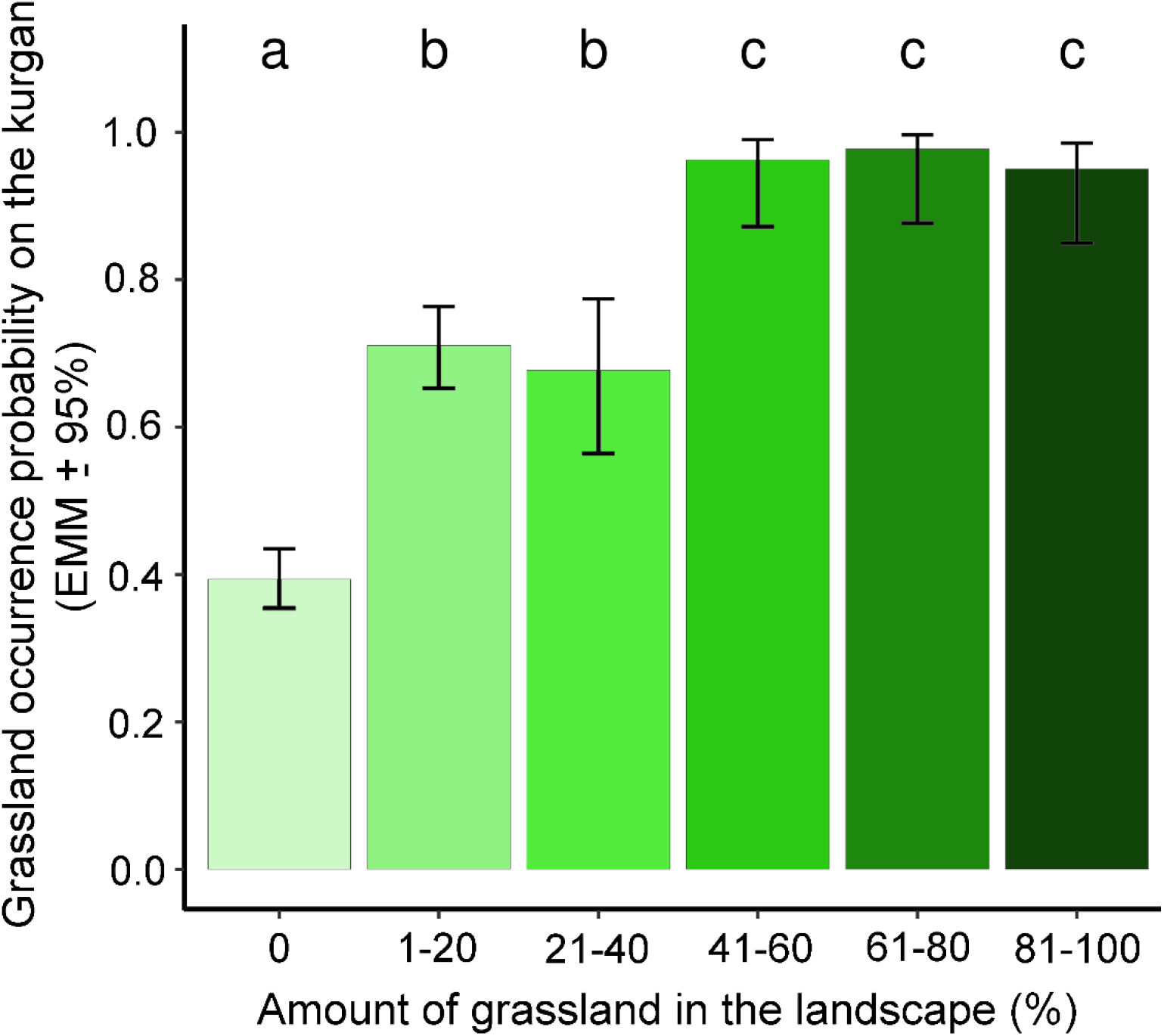
Probability of the occurrence of grasslands on kurgans as a function of grassland amount in the neighbouring landscape (500-m-buffer). Different letters denote significant differences between groups (contrasts from Bayesian logistic generalised regression models; p ≤ 0.05) (n=1032).

We found that kurgans with area-based (77.5%), cultural (66.5%), and combined (73.0%) protection held grasslands with a significantly higher probability compared to non-protected ones (50.8%) (Fig. 3.; Appendices S10-12). Grassland amount in the surrounding landscape was the highest in the case of kurgans with area-based or combined protection. In the case of non-protected kurgans and kurgans with cultural protection, grassland amount in the landscapes was equally low (Fig. 4.; Appendices S13-15).

## Discussion

### Conservation function mediated by the level of landscape transformation

Our study showed that a considerable proportion (58%) of the surveyed kurgans held grasslands and that a majority of them were situated in highly transformed landscapes outside PAs, where they can be considered essential elements of the green infrastructure. The loss of grassland habitats in landscapes around the kurgans was the largest in Ukraine and Russia (Fig. 3). In these countries the large-scale transformation of steppe grasslands to arable land was initiated in the 18^th^ century and accelerated from the 1920’ onwards with the development of agricultural machinery (Smelansky et al., 2012). In the whole Soviet Union approximately 452,000 km^2^ of grasslands had been converted into arable land between 1954 and 1963 during the ‘Virgin Land Campaign’, which also affected large steppe areas in Kazakhstan (Kamp et al., 2016; Smelansky et al.,2012). The grassland amount in the landscape was also low in Hungarian and Serbian study sites, where the considerable loss of dry grassland habitats also dates back to the 18^th^ century (Biró et al., 2018; Deák et al., 2021b).

We found that even in heavily transformed landscapes (grassland amount <1%), kurgans had the potential to preserve grasslands with a high probability (39%) (Fig. 5). This suggests that in such landscapes, kurgans holding grasslands can function as habitat islands, and in this sense, their function is similar to that of other small natural features such as road verges, ravines, forest fringes, hedgerows, mid-field islets, old trees and inselbergs, which occur within and outside the steppe and forest steppe biomes (Hunter et al., 2017). As shown by former studies, the conservation potential of isolated kurgans is supported by their millennia-long existence and relatively undisturbed status, which enables them able to effectively preserve remnants of the original biota despite the marked level of land transformation in the neighbouring landscape (Deák et al., 2020, Dembicz et al. 2020). However, it should be noted here that the number of kurgans reported in the EKDB might show a slight bias towards grassland-covered kurgans, as during data collection some surveyors might favor kurgans that are spectacular in some way, resulting in the overrepresentation of kurgans covered by grasslands.

In landscapes where grassland amount ranges between 1 to 40% kurgans holding grasslands can be essential integral parts of the landscape-level grassland network. By acting as stepping stones, these kurgans can enhance the functional spatial connectivity between the remaining grassland fragments and contribute to the maintenance of meta-population connections of grassland species (Deák et al., 2020).

Landscapes with a grassland share of more than 40% are typical for protected areas of many European regions and also for some Asian countries (such as Mongolia and Kazakhstan), where the majority of the world’s remaining near-natural temperate grasslands can be found (Kamp et al., 2016). In Kazakhstan and Mongolia, all kurgans were covered by grasslands, except those that were built from stones. Based on the findings of previous researches, in these landscapes, the conservation importance of kurgans is rather linked to their high biodiversity and unique flora. Recent studies from Europe (Deák et al., 2021a; Lisetskii et al., 2016,) and Central Asia (Deák et al., 2017) revealed that due to their specific environmental conditions, kurgans often harbour a high number of grassland species that are underrepresented in the neighbouring plain landscapes. Typical examples are the occurrence of forest-steppe plants on the mild north-facing slopes of kurgans situated in the steppe biome (Deák et al., 2017) and steppe plants situated on the top and on south-facing slopes of the kurgans within the forest-steppe biome (Deák et al., 2021a). In this sense, the function of the kurgans is analogous to the patterns described from landmarks, such as inselbergs and dolines, which are also characterized by a high level of topographical heterogeneity (Bátori et al., 2021; Ottaviani et al., 2016).

### Kurgans complement PAs in grassland conservation

We found that kurgans with area-based and combined protection held grasslands with a significantly higher probability compared to non-protected ones (Fig. 3). Our results confirmed that kurgans situated inside PAs could highly benefit from area-based protection even if it was not focused on the kurgan itself. The model predicted the highest probability of grassland occurrence in PAs. It can be considered a promising, but quite evident pattern since protected areas were designated and managed for preserving landscapes that primarily held a high proportion of natural or semi-natural habitats. In such intact landscapes kurgans were less affected by unfavourable anthropogenic disturbances even before they became an integral part of the PA.

Grassland amount in the surrounding landscape was the highest in the case of kurgans with area-based or combined protection, which suggests that the favourable landscape context also provided higher chances for grassland occurrence on kurgans inside PAs (Fig. 4). After the designation of the PAs, kurgans could likely benefit from the area-based protection provided by extensive management measures and prohibition of land transformation actions on the landscape level. Although the direct involvement of kurgans into the system of PAs could likely increase the chance for grassland maintenance, the small size and often isolated location of the mounds makes it a challenging task. Furthermore, as the Hungarian experience shows, legal protection of individual mounds can be a promising tool to overcome this problem, but has a moderate efficiency mostly because it is rather challenging to detect and to mitigate harmful processes in dispersed small objects (Tóth et al., 2018).

Kurgans outside PAs had a high potential for preserving grasslands. This is especially important as kurgans outside PAs (n=846) outnumbered those inside (n=186) by far. In countries situated in the western part of the steppe biome, non-protected kurgans were typically embedded in agricultural landscapes with a low grassland amount. We found that in Hungary, where kurgans are considered “protected landscape elements”, the involvement of kurgans in the regulations connected to the Nature Conservation Law and the single area payment scheme (SAPS) resulted in a slightly higher probability of grassland occurrence compared to the neighbouring Central-European countries. However, as the primary aim of regulations connected to the SAPS is to suppress arable farming on kurgans, it is rather a tool for preventing soil erosion and providing good environmental conditions for spontaneous succession than a measure of active grassland recovery.

Our results suggest that despite the enormous extent of land transformation in the landscape, even non-protected kurgans could effectively preserve grasslands. The practical explanation for this is that ploughing on kurgans with steep slopes can be difficult even with modern agricultural machinery and was even more challenging in the ages of large landscape transformation with less effective agricultural tools (Deák et al., 2016). The protective effect of steep slopes is especially apparent in Bulgaria, where mounds generally have extremely steep slopes, which make them highly unsuitable for arable farming (Apostolova et al., 2022). In this sense, kurgans show similarities with other landscape features such as rocky outcrops and ravines that could escape the plough due to their special physical attributes (Dembicz et al. 2020; Hunter et al., 2017).

### Conservation relevance of cultural values

We found that a considerable proportion of kurgans held various historical and cultural values such as historical buildings, statues, memorial places, medieval roads, border marks and immaterial values (altogether 57 types of cultural values; Appendices S16-17). Since kurgans were built millennia ago, most of the ancient sacred objects originally placed on them have already disappeared. However, as shown by our data, ancient stone pillars and statues are still visible on kurgans in some sparsely populated remote areas in Kazakhstan and Mongolia. Although the builders’ ancient faith has mostly disappeared, subsequent cultures realized the sacredness bound to the kurgans, which resulted in the continuation of its sacred function (Deák et al., 2019b). As shown by our data, the original burial function of kurgans often persists until today. Accordingly, many kurgans still hold existing cemeteries, with graves dating back to medieval times. In Hungary and Serbia, as indicated by numerous sacral buildings and objects, the Christian church had a very strong relatedness to the kurgans, which have served as sacred focal points in the landscape since historical times. In these countries kurgans have often been used as foundation for crosses, calvaries, chapels and even churches since the 10^th^ century, when Christianity became state religion. Although the civilizations that built the kurgans disappeared millennia ago, the sacred heritage represented by the ancient burial mounds has been recognized and respected by the subsequent generations (Deák et al., 2019b).

We found that outside PAs cultural protection could almost double the chance for grassland presence compared to kurgans without cultural values in the same landscape (Fig. 3). This pattern is especially important as kurgans with cultural protection are generally situated in heavily transformed, densely populated anthropogenic lowland landscapes, where grassland habitats are critically endangered (and 13% of kurgans outside PAs hold cultural values). In this respect, kurgans show similarity with sacred natural sites (SNSs) such as church yards, shrines or old cemeteries, which can also preserve parts of the former natural habitats in landscapes under a high anthropogenic pressure (Löki et al.; 2019, Kowarik et al., 2016; Zannini et al., 2021).

Land transformation activities such as ploughing, and afforestation were less typical on kurgans with cultural protection due to their diverse social functions and the respect of the local populations. In order to maintain the societal functions and provide a well-kept appearance of such kurgans, continuous extensive management is often provided by mowing and elimination of woody vegetation, which can also benefit grassland maintenance (Deák et al.; 2020b). This phenomenon shows considerable similarities with the management of SNSs (sacred grooves, saint mountains, shrines, graveyards or church gardens), which can be found across the Globe and are traditionally extensively managed by the local populations for non-production purposes (Zannini et al., 2021).

### Conclusions and conservation outlook

Our continental-scale synthesis revealed that ancient burial mounds, which are the most widespread manmade landmarks in the Eurasian steppes, play a considerable role in grassland conservation. Previous studies found that kurgans can function as safe havens, stepping stones or biodiversity hotspots depending on the landscape context (Deák et al., 2020; Dembicz et al., 2020). We could upscale these findings to a larger geographical scale, as we found that grassland-covered kurgans occur in various scenarios along a landscape transformation gradient across Eurasia. This suggests that kurgans can provide diverse ecological functions mediated by their landscape context.

We found that both area-based protection and the presence of cultural values considerably increased the probability of grassland presence on kurgans. As shown by our results, only a small proportion of kurgans are embedded in PAs. These kurgans indirectly benefit from the extensive land use typical to areas officially designated for nature conservation, but this kind of protection is quite recent and generally not focused on the kurgans. The presence of cultural values considerably contributes to the long-term maintenance of grassland habitats on kurgans. The respect of the local communities provides strong non-official protection, which is based on a social consensus and traditions aiming to preserve the cultural values related to the kurgans. Benefits provided by cultural protection might be especially important as – according to the EKDB – cultural values were most typical in transformed anthropogenic landscapes in the western part of Eurasia.

By holding the last remnants of grasslands, kurgans situated in agricultural or peri-urban areas can serve as additional pillars of grassland conservation. This is especially important, because in fertile lowland landscapes, PAs are generally small, overlap rarely with the distribution areas of many rare or protected species and do not provide a functional connection between their meta-populations (Baranazelli et al., 2022). Taking into account the climate changes expected for the next few decades, we can conclude that a landscape-level network of semi-natural habitats will be of increasingly high conservation importance in the near future (Kirschner et al., 2020). Such a green infrastructure network can allow for an adaptive spatial movement of climate-sensitive species and thus contribute to maintaining their populations. Additionally, kurgans characterized by a variety of climatically different micro-sites can also buffer the effect of changing climate on the site level since climate-sensitive species can easily relocate into microsites matching their environmental needs (Deák et al., 2021a).

Besides those with area-based or cultural protection, there is still a huge proportion of kurgans that does not benefit from any kind of protection. As in the western part of the steppes most kurgans are embedded in arable fields (Apostolova et al., 2022; Deák et al., 2016; Dembicz et al., 2020), including kurgan protection in the system of agri-environmental subsidies may offer a feasible solution for the maintenance or restoration of grasslands on the mounds. The current SAPS-based kurgan protection framework applied in Hungary could be taken as an example for this. However, it should be noted that these regulations predominantly focus on the prohibition of intensive land use on the kurgans and do not include active and targeted measures for supporting grassland restoration on formerly ploughed or degraded kurgans, which would be necessary to restore their ecosystem functions and grassland biodiversity. Considering related costs in CAP subsidies (e.g. in form of additional support for voluntarily performed active grassland restoration) would be a nature-based solution for the restoration of kurgan-related ecosystem services such as pollination, weed suppression, pest control and increased landscape value. It would be possible to set up a general SAPS-based framework for kurgan protection, which might be adapted to the specific conditions of each country. This framework could be flexibly adapted to non-EU countries or West-European countries, where other types of prehistoric burial mounds occur.

The EU Biodiversity Strategy has the ambition of increasing the area of non-cultivated high-diversity landscape features (HDLFs, e.g. hedgerows, flower strips and riparian corridors) to at least 10% of the utilized agricultural area by 2030. Grasslands on kurgans could also be recognized as HDLFs, forming an additional pillar for their conservation and restoration. Besides the SAPS-based framework, kurgan protection might also be achieved with bottom-up public participation since the mounds are highly recognized by local populations and can thus be proper targets of non-governmental organizations focusing on the preservation of local cultural and natural heritage (Valkó et al., 2018).

Recently a growing number of studies found that the global system of protected areas does not provide a sufficient level of biodiversity protection (Wauchope et al., 2022). The effectiveness of the global PA system can be increased to some extent by designating new PAs using improved spatial planning tools or by increasing the efficiency of protection within PAs. However, these measures all have limits as the land for PAs is also demanded by agriculture, industry and infrastructural development. Our study highlights that to complement and support the system of PAs, it is crucial to acknowledge the conservation potential of sites that – thanks to their associated cultural values – can harbour natural habitats even in non-protected landscapes. Our results suggest that an integrative socio-ecological approach in conservation could support the positive synergistic effects of conservational, landscape and cultural values.

## Acknowledgements

We are thankful to G. Batdelger, M. Biró, Z. Gavrilović, D. Grabovác, R. Mirić, Z. Molnár, Z. Nagygyörgy, S. Spremo, P.I. Tóth, R. Vesović and the entire EKDB community for their help in exploring kurgans. This study was supported by the Hungarian National Research, Development and Innovation Office (NKFI FK 135329). The work of OV and BD was supported by the Hungarian National Research, Development and Innovation Office (NKFI KKP 144096). PB, RG and BD were supported by the Hungarian National Research, Development and Innovation Office (NKFIH KKP 133839). IA and GN were supported by the Bulgarian National Science Fund (contract KП-06-H21/2, 2018). RG was supported by the János Bolyai Research Scholarship of the Hungarian Academy of Sciences. ID was supported by the Polish National Science Centre (grant no. DEC-2013/09/N/NZ8/03234). BS and IM were supported by the Scientific Research Committee, Poland (grants no. 2PO4G04627 and no. NN304081835). SB was supported by the NKFIH KDP 967901 grant. The authors are grateful to Aiko Huckauf for the linguistic edition of the manuscript.

## Author contributions

The structure of the online database was designed by BD, ÁB, FB, CT, MB, and OV. The online database was programmed and maintained by MB. Field data collection was done by BD, ÁB, ID, IA, PB, RG, CT, JD, IM, BS, GN, FL, ZB, SK, SB, LG, TB, IS, ÁM, and OV. Data preparation for the analyses was done by BD, ÁB, and FB. Data analyses were performed by ZR and BD. The conception of the manuscript was compiled by BD and OV. The first draft of the manuscript was written by BD and OV, and was revised by ZR, PB, RG, ID and IA. All authors commented on previous versions of the manuscript. All authors read and approved the final manuscript.

## Data availability

The data that support the findings of this study are available in figshare with the identifier https://figshare.com/s/9c168f820745187d2b8c.

## List of Appendices

**Appendix S1**. Number of kurgans holding grasslands across the six studied countries (n=1032).

**Appendix S2**. Probabilities calculated for grassland occurrence on kurgans situated in the six studied countries (Bayesian logistic generalised regression models).

**Appendix S3**. Contrasts and odds ratios calculated for grassland occurrence on kurgans situated in the six studied countries (Bayesian logistic generalised regression models). P-values were adjusted for multiple comparisons with Bonferroni’s method. Significant effects are marked with boldface.

**Appendix S4**. Number of kurgans situated in landscapes with different amount of grasslands (within a 500 m buffer around the kurgans) (n=1032).

**Appendix S5**. Estimated marginal means (EMM) calculated for grassland amount in the landscape surrounding the kurgans (within a 500 m buffer) situated in the six studied countries (proportional odds logistic regression models).

**Appendix S6**. Contrasts and estimates calculated for grassland amount in the landscape surrounding the kurgans (within a 500 m buffer) in the six studied countries (proportional odds logistic regression models). P-values were adjusted for multiple comparisons with Bonferroni’s method. Significant effects are marked with boldface.

**Appendix S7**. Number of kurgans holding grasslands and ordinal values of grassland amount in the neighbouring landscape (n=1032).

**Appendix S8**. Probabilities calculated for grassland occurrence on kurgans as a function of grassland amount in the landscape surrounding the kurgans (within a 500 m buffer) (proportional odds logistic regression models). Significant effects are marked with boldface.

**Appendix S9**. Contrasts and odds ratios calculated for occurrence of grasslands on kurgans as a function of grassland amount in the landscape surrounding the kurgans (within a 500 m buffer) (proportional odds logistic regression model). P-values were adjusted for multiple comparisons with Bonferroni’s method. Significant effects are marked with boldface.

**Appendix S10**. Number of kurgans holding grasslands across the four protection types (n=1032).

**Appendix S11**. Probabilities calculated for grassland occurrence on kurgans under different types of protection (Bayesian logistic generalised regression models).

**Appendix S12**. Contrasts and odds ratios calculated for grassland occurrence on kurgans under different types of protection (Bayesian logistic generalised regression models). P-values were adjusted for multiple comparisons with Bonferroni’s method. Significant effects are marked with boldface.

**Appendix S13**. Number of kurgans under different type of protection that hold grasslands in landscapes with various amount of grasslands (n=1032).

**Appendix S14**. Estimated marginal means (EMM) calculated for grassland amounts in the landscape surrounding kurgans that are under different level of protection (proportional odds logistic regression model). Significant effects are marked with boldface.

**Appendix S15**. Contrasts and estimates calculated for grassland amount in the landscape surrounding the kurgans (within a 500 m buffer) under different type of protection (proportional odds logistic regression models). P-values were adjusted for multiple comparisons with Bonferroni’s method. Significant effects are marked with boldface.

**Appendix S16**. Cultural and historical objects on mounds covered by grasslands.

**Appendix S17**. Number of occurrences of historical, cultural and sacred values confined to the kurgans (n=1072).

